# GENOMIC ANALYSIS OF THE STREPTOMYCES SP. LV42-5 ISOLATED FROM INDUSTRIAL MINE DUMPS IN SHEPTYTSKYI

**DOI:** 10.64898/2025.12.04.691995

**Authors:** I. Roman, O. Makar, O. Stasyk, V. Fedorenko, O. Gromyko

## Abstract

Actinomycetes, particularly within the genus *Streptomyces*, remain the most prolific bacterial source of secondary metabolites for medical and biotechnological applications. However, contemporary drug discovery is increasingly challenged by the high rate of rediscovering known molecules and the difficulty of activating “silent” biosynthetic gene clusters (BGCs) under standard laboratory conditions. To address this, bioprospecting in extreme environments, such as heavy metal-contaminated mine dumps, has emerged as a strategic approach to uncover strains with unique metabolic adaptations and chemically diverse natural products. This study reports the whole-genome sequencing, assembly, and comprehensive bioinformatic analysis of Streptomyces sp. Lv42-5, an extremotolerant strain isolated from the rhizosphere of birch trees (*Betula pendula*) growing on an industrial mine dump in Sheptytskyi, Ukraine. Genomic sequencing utilizing the Illumina platform followed by de novo assembly yielded a high-quality draft genome of 9.84 Mbp with a G+C content of 71%. Phylogenomic analysis using the GTDB and ANI calculation revealed that strain Lv42-5 shares only 93.36% ANI with its closest relative, *Streptomyces diastatochromogenes*. This value falls well below the 95–96% species delineation threshold, confirming Lv42-5 as a taxonomically novel species. Functional annotation via the RAST server indicated a genome heavily dedicated to metabolic processes, particularly amino acid and carbohydrate metabolism, while lacking genes for motility and photosynthesis. Crucially, the genome encodes a robust genetic arsenal for heavy metal resistance, including specific mechanisms for tolerating copper, cobalt, zinc, and cadmium, reflecting the strain’s successful adaptation to its metalliferous habitat. Genome mining using antiSMASH 8.1 uncovered a rich biosynthetic landscape comprising 43 putative gene clusters. These findings establish *Streptomyces* sp. Lv42-5 as a novel, stress-adapted species with significant dual potential for bioremediation of heavy metal pollutants and the discovery of novel therapeutic agents.

---

Actinomycetes, particularly members of the genus *Streptomyces*, are a group of prokaryotes well-known for their production of secondary metabolites. This remarkable biosynthetic capacity stems from the significant portion of their genomes, often 5–20 % in Streptomyces, dedicated to biosynthetic gene clusters (BGCs), making them the most productive bacterial source of such compounds for medical and biotechnological applications [10]. While over 7,600 compounds have been identified from actinomycetes, contemporary drug discovery efforts are frequently hampered by the high rate of rediscovery of known molecules. A further challenge lies in exploiting their full genomic potential, as a substantial number of their BGCs remain “silent” or are expressed at insignificant levels under standard laboratory conditions [21]. To circumvent these obstacles and tap into novel chemical diversity, researchers are increasingly focusing on bioprospecting for actinomycetes in underexplored and extreme environments, including those contaminated with various pollutants, which may harbor strains with unique and previously uncharacterized biosynthetic pathways.

Actinomycetes isolated from environments contaminated with heavy metals demonstrate significant potential for both bioremediation and the discovery of novel bioactive compounds [7]. These microorganisms, particularly from the genus Streptomyces, can tolerate and often accumulate toxic metals like cadmium (Cd), lead (Pb), and nickel (Ni), making them excellent candidates for cleaning up polluted sites [13]. An aspect of these extremophilic bacteria is how environmental stress can unlock their biosynthetic potential. The presence of heavy metals can trigger the expression of silent biosynthetic gene clusters, leading to the production of unique secondary metabolites. This was observed in *Streptomyces* sp. WU20, which synthesized a novel antibiotic specifically when exposed to nickel stress [20]. That can be discovered from bacteria thriving under harsh conditions. This strategy of “environmental stress-induced activation” allows researchers to tap into the hidden genomic potential of these microorganisms.

Mine dumps, characterized by their harsh physicochemical conditions and high concentrations of heavy metals, are increasingly targeted as valuable ecosystems for isolating actinomycetes with unique metabolic and biosynthetic capabilities. The extreme selective pressures in these environments foster the evolution of microorganisms that not only exhibit remarkable resistance to heavy metals but also possess the capacity to produce novel bioactive secondary metabolites. This potential has been demonstrated by *Streptomyces* strains from Moroccan mining soils, which displayed significant antimicrobial activity that could be modulated by modifying culture conditions [1]. Furthermore, these sites are hotspots for taxonomic novelty, leading to the discovery of new species such as *Streptomyces manganisoli, Actinorectispora metalli, Amycolatopsis saalfeldensis*, and *Catellatospora koreensis* [5-6, 11-12]. Bioprospecting in such locations has also yielded tangible therapeutic leads, with actinomycetes from waste dumps producing previously uncharacterized antifungal and antibacterial compounds active against drug-resistant pathogens [18].

This study reports the whole-genome sequencing and comprehensive annotation of an actinomycete strain isolated from the rhizosphere of birch trees (*Betula pendula*) inhabiting industrial mine dumps in Sheptytskyi (formerly Chervonohrad), Ukraine [14]. The primary objective of this investigation was to characterize the complete genome of this extremotolerant isolate, with a specific focus on identifying and analyzing its biosynthetic gene clusters (BGCs) to uncover its potential for producing novel secondary metabolites.

## Materials and Methods

### Bacterial Strain and Genomic Data

The actinomycete strain *Streptomyces* sp. Lv42-5, previously isolated from the rhizosphere of birch trees growing on heavy metal-contaminated mine dumps in Sheptytskyi, Ukraine, was the subject of this study [14]. Genomic DNA was extracted, and the whole genome was sequenced using the Illumina platform. A *de novo* assembly of the raw sequencing reads was subsequently performed to generate a draft genome.

### Bioinformatic Analysis

The quality of the assembled draft genome was evaluated for completeness and contamination using CheckM [15]. Taxonomic assignment was conducted using the Genome Taxonomy Database Toolkit (GTDB-Tk). To refine the species-level classification, the Average Nucleotide Identity (ANI) was calculated against the closest relative, *S. diastatochromogenes*. Comprehensive genome annotation was performed using the Bakta pipeline [19] to identify coding sequences (CDS), tRNAs, rRNAs, and other genetic features. The predicted protein-coding genes were then functionally categorized into subsystems with the Rapid Annotation using Subsystem Technology (RAST) server [2]. To explore the strain’s biosynthetic potential, the genome was mined for secondary metabolite biosynthetic gene clusters (BGCs) using the antiSMASH platform (version 8.1). BGCs of interest were further analyzed by comparing them against the Minimum Information about a Biosynthetic Gene cluster (MIBiG) database to identify homologous clusters with known products. Raw genomic data are available from the authors upon request.

## Results and Discussion

*De novo* assembly of the genome of strain Lv42-5 resulted in a draft genome of 9,842,483 bp with a GC content of 71 %, distributed across 102 contigs. Quality assessment indicated 100 % completeness and 0.95 % contamination, with an N50 value of 192,601 bp. Taxonomic analysis using the Genome Taxonomy Database (GTDB) revealed the closest relative to be *Streptomyces diastatochromogenes*, with an Average Nucleotide Identity (ANI) of 93.36 %. This ANI value is below the conventional species delineation threshold of 95–96 %, strongly suggesting that strain Lv42-5 represents a novel species within the genus Streptomyces. Genomic annotation using Bakta identified 8,923 coding sequences (CDS), 90 tRNAs, 5 rRNAs, 29 non-coding RNAs (ncRNAs), and one transfer-messenger RNA (tmRNA). Functional analysis of the CDS revealed a genome predominantly dedicated to metabolic processes. The largest categories were associated with the metabolism of amino acids and derivatives (502 CDSs), carbohydrates (444 CDSs), cofactors and vitamins (255 CDSs), and proteins (241 CDSs). Other significant metabolic functions included fatty acid and lipid metabolism (206 CDSs), respiration (144 CDSs), and nucleotide metabolism (138 CDSs). CDSs related to stress response (80), cell wall biogenesis (62), virulence and defence (58), and membrane transport (53) were also well-represented. In contrast, genes associated with secondary metabolism (14), dormancy and sporulation (13), and mobile elements (2) were less abundant. Notably, no genes were detected for photosynthesis, motility, or cell division. (fig 1).

**Fig. 1.**
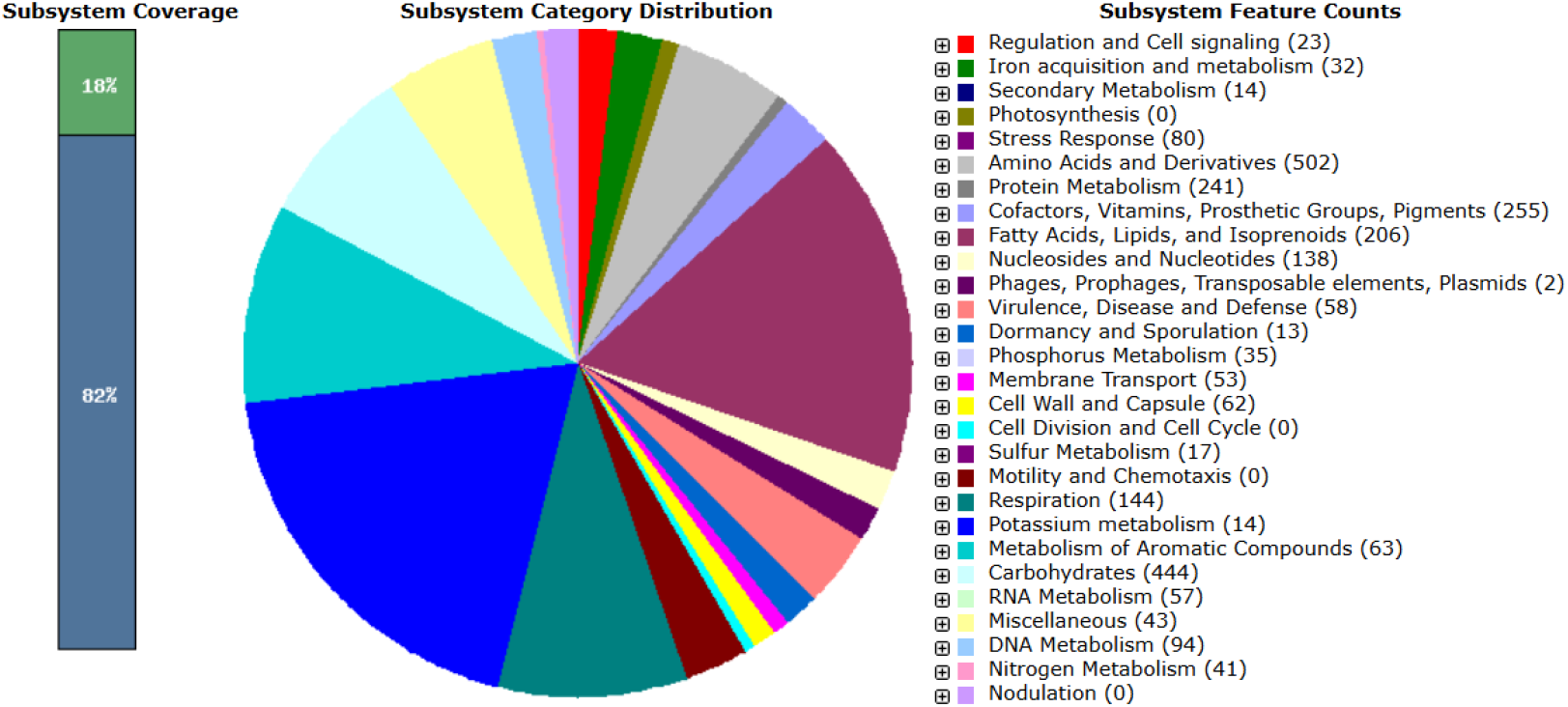
RAST annotation summary of strain Lv42-5. The RAST annotation robot assigns names and functions to protein-coding genes via their subsystem technology. The green colour represents features that are found in RAST subsystem. The blue colour represents features not assigned to a subsystem

Of particular ecological relevance, the genomic analysis identified a substantial number of genes conferring resistance to heavy metals such as copper (Cu), cobalt (Co), zinc (Zn), and cadmium (Cd). This genetic arsenal is a key adaptation, providing the necessary mechanisms for survival and proliferation within the toxic, metal-laden environment of the mine dumps from which the strain was isolated [9].

To investigate the strain’s biosynthetic potential, particularly in relation to its adaptation to the metal-rich environment, the genome was analyzed using the antiSMASH platform. This analysis revealed the presence of 43 biosynthetic gene clusters (BGCs). Comparative analysis of the biosynthetic gene clusters revealed a wide spectrum of novelty. Among the identified BGCs, 12 exhibited high homology to known pathways, three showed medium homology, and 11 displayed low homologies. Significantly, 17 BGCs had no detectable homology to any characterized clusters in public databases, representing a substantial reservoir of potentially novel biosynthetic pathways. The highly homologous clusters included those responsible for the synthesis of ε-poly-L-lysine, geosmin, ectoine, hopene, and various spore pigments. These BGCs are commonly conserved among actinomycetes, as their products fulfil fundamental functions essential for cell viability, stress tolerance, and development. While the majority of these highly conserved BGCs were predicted to synthesize peptides of uncharacterized function, one BGC showed high sequence homology to the known biosynthetic pathway for largimycins, a class of non-ribosomal peptides (NRPs) recognized for their potent cytotoxic and anti-cancer activities [3]. Among the highly homologous BGCs, a cluster responsible for the synthesis of albaflavenone, a known terpene antibiotic, was also identified [22], an antibiotic from this family was also identified in our previous work on actinomycetes from the Crimean Peninsula [17]. Furthermore, the genomic analysis identified biosynthetic gene cluster that exhibited significant homology to the simocyclinone cluster. This antibiotic leads to the inhibition of bacterial DNA gyrase, an essential enzyme for bacterial replication [4]. However, the biosynthetic gene clusters exhibiting low or no homology to characterized compounds need the most significant attention. These clusters represent a compelling source for natural product discovery, as they are predicted to direct the synthesis of novel compounds possessing unique chemical scaffolds and biological activities.

A significant emphasis was placed on three clusters predicted to synthesize siderophores, which are high-affinity iron-chelating compounds that can also sequester other metals. Among these, Cluster 16.1 was of particular interest, as it exhibited high sequence homology to the well-characterized BGCs for desferrioxamine in *Streptomyces argillaceus* and *Streptomyces coelicolor* A3(2), as well as to the legonoxamine A BGC from *Streptomyces* sp. MA37 (fig. 2). Siderophores belonging to this hydroxamate class, such as desferrioxamine B, are widespread among streptomycetes and are recognized for their potential use in bioremediation applications due to their strong metal-binding capabilities [8]. In contrast to the desferrioxamine-like cluster, the other two putative siderophore BGCs (13.3 and 70.1) lacked significant homology to any characterized clusters in public databases. This finding suggests that these gene clusters may direct the synthesis of novel siderophores with previously undescribed chemical structures, highlighting the strain’s potential for producing unique metal-chelating agents.

**Fig. 2.**
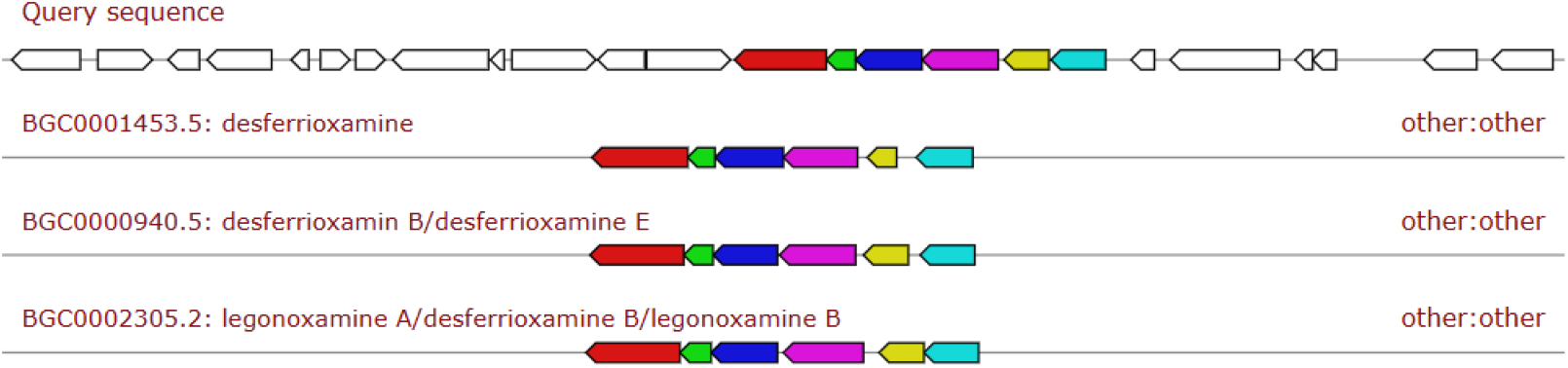
Comparative analysis of the putative siderophore biosynthetic gene cluster 16.1 of the strain Lv 42-5 showing similarity to known clusters encoding desferrioxamine and legonoxamine biosynthesis (BGC0001453.5, BGC0000940.5, and BGC0002305.2) from the MIBiG database

Genomic analysis also identified a melanin biosynthetic gene cluster (BGC), which included a tyrosinase gene exhibiting high homology to its counterpart in *Streptomyces avermitilis*. The resulting melanin polymer is rich in functional groups, such as carboxyl, hydroxyl, and phenolic units, that provide numerous negatively charged sites for chelating metal cations. This molecular architecture enables the strain to sequester and immobilize toxic heavy metals like cadmium and lead, thereby reducing their bioavailability and mitigating cellular damage. This detoxification mechanism operates both on the cell surface and within the cell wall, creating a robust protective barrier that effectively limits the influx of harmful ions into the cytoplasm [16].

In conclusion, the comprehensive genomic analysis of *Streptomyces sp*. Lv42-5 confirms its status as a novel species uniquely adapted to the harsh, metal-contaminated environment of mine dumps. Its genome is equipped with a robust arsenal of heavy metal resistance genes and a diverse collection of biosynthetic gene clusters responsible for producing metal-chelating compounds like siderophores and melanin. The identification of BGCs for both known and potentially novel siderophores highlights this strain as a promising candidate not only for bioremediation applications but also as a valuable source for the discovery of new natural products. This work underscores the potential of bioprospecting in extreme environments to uncover microorganisms with unique metabolic and biosynthetic capabilities.

## Acknowledgements

The authors would like to thank the Ministry of Education and Science of Ukraine for its grant support (BG-18F).

